# The cryo-EM structure of mouse radial spoke 3 reveals a unique metabolic and regulatory hub in cilia

**DOI:** 10.1101/2025.05.19.654877

**Authors:** Yanhe Zhao, Kangkang Song, Amirrasoul Tavakoli, Long Gui, Angeles Fernandez-Gonzalez, Song Zhang, Petras P Dzeja, S. Alex Mitsialis, Xuewu Zhang, Daniela Nicastro

**Affiliations:** Department of Cell Biology, University of Texas Southwestern Medical Center, Dallas, Texas 75235; Division of Newborn Medicine, Department of Pediatrics, Boston Children’s Hospital, Harvard Medical School, Boston, MA, USA; Department of Cardiovascular Medicine, Mayo Clinic, Rochester, MN 55905; Department of Pharmacology, University of Texas Southwestern Medical Center, Dallas, Texas 75390; Department of Biophysics, University of Texas Southwestern Medical Center, Dallas, Texas 75390

## Abstract

Cilia are complex, microtubule-based organelles that protrude from many eukaryotic cells and have important roles in sensing, signaling, and motility. Recent studies have revealed the atomic structures of many multi-component ciliary complexes, providing new insights into their mechanisms of action that are vital for cilia’s biological functions. However, little is known about the structure, proteome, and function of full-length radial spoke 3 (RS3), which is distinct from the structurally well-characterized RS1 and RS2. Radial spokes are conserved megadalton complexes that transmit mechanochemical signals from the central pair of microtubules to the dynein motors, thereby coordinating ciliary motility. Here, we combined cryo-electron microscopic single-particle reconstruction, cryo-electron tomography (cryo-ET), proteomic analysis, and computational modeling to determine the 3D structure and atomic model of RS3 from mouse respiratory cilia. Our structure reveals all protein components of RS3, including regulatory and metabolic enzymes, such as a protein kinase A subunit, adenylate kinases and malate dehydrogenases. We have confirmed the important role of adenylate kinase 7 in RS3 by cryo-ET analyses of respiratory cilia in AK7-deficient mice, which display primary ciliary dyskinesia. Our findings suggest that RS3 is an important regulatory hub and cluster of metabolic proteins that helps to maintain ATP at the levels required for sustained dynein motor activity and ciliary beating. This work advances our understanding of the structure and function of RS3 in ciliary motility and provides insights into the etiology of ciliopathies.

## Introduction

Cilia are hair-like structures that protrude from the surface of most eukaryotic cells. Motile cilia undulate rapidly, displacing surrounding fluids to cause either flow over a tissue or motility through the fluid, whereas non-motile cilia are sensory organelles. Cilia are essential for human health, because impaired ciliary assembly or motility causes ciliopathies, such as primary ciliary dyskinesia (PCD), which is characterized by infertility, chronic respiratory issues, misplaced organs, and excess cerebrospinal fluid (hydrocephalus)^1–3^. Decades of research have provided many insights into the structure and function of motile cilia, yet precisely how the thousands of dynein motors are spatio-temporally regulated to generate the waveforms typical of beating cilia and flagella remains incompletely understood.

From protozoa to mammalian cells, motile cilia contain a microtubule-based scaffold called the axoneme that usually consists of a cylinder of nine doublet microtubules (DMTs) surrounding two central singlet microtubules (central pair complex, CPC); this organization is commonly referred to as [9+2] (Fig. 1a). Hundreds of highly conserved ciliary proteins^4^ form macromolecular superstructures that are arranged in 96 nm repeating units along the axoneme, including the outer and inner dynein arms (ODAs and IDAs), which drive motility, and regulatory complexes like the nexin-dynein regulatory complex (N-DRC), the calmodulin and spoke associated complex (CSC), and a triplet of radial spokes (RSs) (Fig. 1b-c). Cryo-electron microscopy and tomography (cryo-EM/ET) studies have characterized the 3D structures and subunit organizations of most of these ciliary complexes^5–13^. Recent cryo-EM single particle reconstructions of ciliary complexes have allowed determining their atomic models^14–17^. A notable exception, however, is the full-length third RS (RS3), for which the distal structure and protein composition have remained elusive^16,18^.

**Fig. 1.**
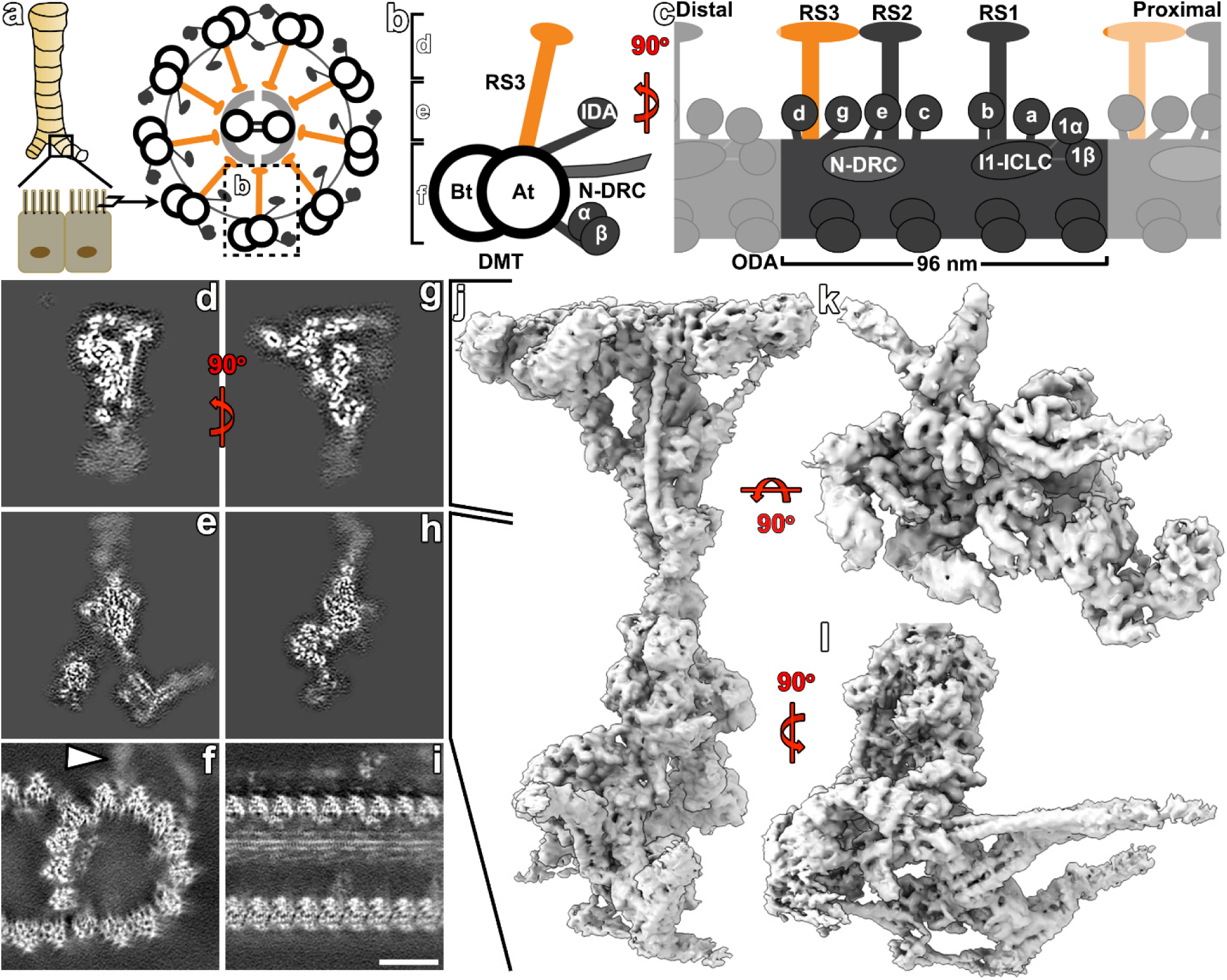
Structural organization of mouse respiratory cilia and cryo-EM single particle reconstruction of the full-length radial spoke 3 (RS3). **a)** Cartoons show that mouse tracheal epithelial cells are multi-ciliated and contain a typical [9+2] axoneme (viewed from distal tip to proximal base of the cilium). Dotted box highlights one doublet microtubule (DMT). **b, c)** Cartoons depict the organization of the 96 nm axonemal repeat in cross-sectional (b) and longitudinal view (c). **d-i)** Slices through the cryo-EM 3D reconstructed RS3 head-neck (d, g) and base regions (e, h), as well as the A-tubule of the DMT (f, i) in cross-sectional (d-f) and longitudinal view (g-i). Note that the blurred densities in the overlap regions between the three reconstructions imply flexibility of these regions relative to each other (arrowhead in f indicates base of RS3). **j-l)** Isosurface renderings of the 3D reconstructed RS3 show the full-length in longitudinal view (j; corresponds to panels g and h), in top-view of the RS3 head (k) and cross-sectional view of the RS3 base (l; corresponds to panel e). Scale bar: (d-i) 10nm.

Mutations in single RS proteins often cause severe motility defects, ciliary paralysis, and ciliopathies^19–25^, so RSs are essential for ciliary motility. RSs span the 40 nm distance between the CPC and the DMT, to which they are attached. They are thought to transduce mechano-chemical signals from the CPC to regulate dynein motor activity and thus ciliary beating^26–28^. However, the mechanisms by which the RS-CPC system transmits regulatory signals to the dyneins is not understood on a molecular level. The overall T-shape of RSs and their spacings of 32 nm, 24 nm, and 40 nm are highly conserved^19^. Previously, all RSs were thought to be identical, but we have demonstrated that RS3 is structurally distinct from RS1 and RS2^29^, differences that are likely to be of functional significance. Mutations that affect RS1 and RS2 but leave RS3 intact caused less severe pathologies in human PCD patients than mutations that alter RS3, highlighting the importance of the latter^19,20^. Previous studies have established the protein composition, 3D structure and atomic model of RS1 and RS2 in several species^16–18,30^, but the unique proteome, structure, and function(s) of RS3 remain poorly characterized. Reasons for this limited characterization include for example, that most previous proteomic analyses of cilia and their isolated complexes used the unicellular algae *Chlamydomonas* as model organism and identified over 600 different, highly conserved proteins^4,31^. Comparison between *Chlamydomonas* wildtype and the presumed “RS-less” *pf14* mutant successfully identified at least 23 RS proteins (RSP1-23), but these were mostly RS1 and RS2 proteins, because *Chlamydomonas* flagella contain only two full-length RSs, RS1 and RS2, and a shorter RS3S (which is identical to the RS3-base), that still assembled in *pf14*^32^.

In this study, we solve the cryo-EM structure of intact, full-length RS3 from mouse respiratory cilia. Resolution of the base-stalk and neck-head reconstructions reach 4.7 Å and 7.1 Å resolution, respectively. By combining these cryo-EM structures with proteomics data and computational modeling, we can identify and situate all protein components in RS3 – adding ten proteins to the four previously known RS3 subunits. Our data provide an atomic model of the unique, full-length RS3 structure. Surprisingly, RS3 contains several proteins important for phospho-regulation (e.g. regulatory subunit of proteins kinase A, and A-kinase anchoring protein), and enzymes involved in energy metabolism, including adenylate kinases AK7 and AK9, and a malate dehydrogenase (MDH) heterodimer. Our findings suggest that RS3 serves both regulatory and metabolic functions by maintaining an appropriate ATP/ADP ratio along the ciliary length, thereby sustaining dynein-mediated ciliary beating.

## Results

### RS3 protein candidates identified by comparative proteomics analysis

By studying the conserved CSC in *Tetrahymena* cilia, which have three full length RSs, we learned that the proteins CFAP61 (RSP19) and CFAP251 (WDR66) localize to the RS3 base^7^. Work in other labs has identified two additional RS3 components: the leucine-rich repeat-containing protein LRRC23 likely in the distal RS3 region^21^ and the CSC-protein CFAP91 that connects the bases of RS2 and RS3^17^. In our previous study, cryo-ET class averages of cilia from a *Tetrahymena* FAP61-knockout strain (FAP61-KO) showed that the RS3 densities were either partially reduced or completely missing (Supplementary Fig. 1a-d)^7^. Mild salt-extraction with 0.6 M NaCl had little effect on RSs in wildtype *Tetrahymena* axonemes (RSs in general are stably attached to the axoneme, requiring potassium iodide treatment for extraction^33,34^), but the remaining RS3s were completely extracted by this treatment from FAP61-KO axonemes, as shown by negative-staining EM (compare Supplementary Fig. 1e,f).

We performed comparative liquid chromatography-mass spectrometry (LC-MS) on these extracts and identified twenty RS3 protein candidates in *Tetrahymena*, in addition to two of the previously identified proteins (Supplementary Table 1). Our proteomic analysis also detected the majority of the *Tetrahymena* homologues of known RS1 and RS2 proteins^30,34,35^, but their abundance was comparable between WT and FAP61-KO (Supplementary Table 2). This supports our previous interpretation that the RS3 proteome is distinct from RS1 and RS2^19,29^. Our previous study using co-IP with CSC proteins identified four adenylate kinase family members in the *Tetrahymena* axoneme, AK1 and AK7-9^7^, but here the WT-FAP61-KO comparison only identified AK7 and AK9 as RS3-specific candidates (Supplementary Table 1). AK7 and AK9 are conserved among ciliated species that have three full-length RSs, such as sea urchin and mammals, but not in *Chlamydomonas*, which has a short RS3S that lacks the distal region. This observation suggests that AK7 and AK9 localize to the distal half of RS3. AK7 and AK9 were also identified in a recent cross-linking mass spectrometry (XL/MS) study of *Tetrahymena* cilia, but they were interpreted as heterodimeric CPC components^36^. Although our data show that this assignment is incorrect, the cross-linking data indicate that AK7 and AK9 interact with each other and with CPC projection protein(s). AK7 and AK9 were also reported as ciliary proteins with unknown roles in a recent human ciliary proteomic study^31^. The *Tetrahymena* proteomic analysis also identified two LRRC family proteins. A BLAST search of mouse LRRC23 against the *Tetrahymena* genome reveals only four hypothetical proteins with relatively low sequence identity. However, AlphaFold 2 structure predictions suggest that one of the here-identified RS3 candidates from the LRRC family (hypothetical protein TTHERM_00105260), is an LRRC23 homolog (previously called LRRC23-like 2^37^).

Adenylate kinases catalyze the transfer of phosphate groups between adenine nucleotides and play important roles in cellular energy metabolism, e.g., maintaining adenine nucleotide homeostasis^38^. Mutations in AK7 have been linked to PCD in mice^39^ and to human infertility^40^. We used cryo-ET of respiratory cilia from WT and AK7-deficient (*AK7^-/-^*) mice to learn whether AK7 is a conserved RS3 component. Respiratory cilia isolated from AK1-knockout (*AK1^-/-^*) mice were analyzed as an additional control; RS3 should be unaffected in this mutant because our *Tetrahymena* proteomic data showed comparable levels of AK1 in salt-extracted WT and FAP61-KO axonemes (Supplementary Table 2). The 3D structures (Supplementary Table 3) revealed that no RS defects were present in cilia from WT and *AK1^-/-^* mice (Supplementary Fig. 1g-h,k-l), but *AK1^-/-^* repeats lacked a previously uncharacterized, rod-shaped structure between RS1 and RS2 (Supplementary Fig. 1i-j,m-n, arrowheads). In contrast, in *AK7^-/-^* mouse cilia, the RS3 head and neck regions were missing, whereas other axonemal structures appeared normal (Supplementary Fig. 1o-r). Classification analysis focused on the remaining RS3 structure in the *AK7^-/-^* axonemal repeat showed three classes with different levels of RS3 truncation (Supplementary Fig. 1s-x, arrowheads), with the shortest class 1 (25% of repeats) resembling RS3S in *Chlamydomonas* flagella (Supplementary Fig. 1s-t).

Given RS3’s consistent truncation in *AK7^-/-^* mice, we conducted a comparative LC-MS analysis using axonemes isolated from WT and *AK7^-/-^*respiratory cilia to identify RS3 protein candidates in mice. The results revealed markedly decreased levels for 10 proteins (mutant/WT ratio <0.1), including AK9 and AK7, which were both identified as RS3 candidates in our *Tetrahymena* proteomic analysis, and LRRC23, a previously identified RS3 component^21^ (Supplementary Fig. 2, red dots). This suggests these proteins are RS3 components and that they strongly depend on AK7 for their stable assembly into RS3, probably through protein-protein interactions. Additional proteins were reduced to varying degrees as shown in Supplementary Figure 2. In contrast, levels of AK1 (located between RS1 and RS2 as shown in Supplementary Fig. 1k-n) and AK8 (previously reported to interact with RSPH4A and RSPH3B in RS1^18^) showed comparable abundance in axonemes from WT and *AK7^-/-^* mice (Supplementary Fig. 2, green dots, and Supplementary Table 4), confirming our *Tetrahymena* RS3 proteomic data (Supplementary Table 2). Overall, our proteomic results provided critical support for modeling the atomic structure of RS3.

### Cryo-EM structure of intact mouse RS3

For the cryo-EM single particle reconstruction of well-preserved, full-length RS3, we isolated axonemes from mouse trachea using an optimized calcium-shock method (see methods), splayed the axonemes into DMTs using ATP and mild salt-treatment (avoiding protease-treatment), and recorded 50,000 cryo-EM images of intact DMTs containing 280,000 axonemal repeats (Supplementary Fig. 3a and Supplementary Table 3, see methods for details). The initial 3D reconstruction of the DMT achieved 4-6 Å resolution but showed blurred density for RS3 (arrowhead in Fig. 1f; arrows in Supplementary Fig. 3e), presumably due to RS3’s flexibility relative to the DMT. After optimizing the image processing workflow, including moving the alignment refinement to two sub-regions of RS3 (Supplementary Figs. 3b, 4a) and selective signal subtraction, the RS3-base/stalk and the RS3-neck/head regions were clearly visible in the 3D reconstructions, achieving resolutions of 4.7 and 7.1 Å, respectively (Supplementary Fig. 4b and Supplementary Table 3). A composite map was generated by combining these two partially overlapping maps, revealing the overall RS3 structure that has a length similar to RS1 and RS2. RS3 consists of a bulky base with two major DMT-attachment sites (i.e., docking factor CFAP57 interacting with CFAP251 (“back-prong”) and the tails of IDA *d* and IDA *g* (“front-prong”), respectively), a slim stalk that might allow some pivotal flexibility of the distal RS3 region relative to its base, and a neck region with four major “struts” supporting the flat-shaped head region that interfaces with the CPC projections (Fig. 1d-e, g-h, j-l).

### Integrated approaches permit an atomic model of mouse RS3

The previously published atomic model of the human RS3 base region (PDB: 8J07) could be readily docked into our RS3 cryo-EM map, however, we “mutated” all included protein sequences from human to the corresponding mouse proteins and performed individual residue refinement to our 3D density during the model building step (see methods for details). Most of the proteins in the base had comparable levels between WT and *AK7^-/-^* mouse axonemes in our proteomic analysis (Supplementary Fig. 2, green dots), consistent with our cryo-ET results showing that the RS3 base region remained intact in the mutant (Supplementary Fig. 1o-p, s-x).

The 7.1 Å cryo-EM map of the RS3 head-neck-stalk regions clearly shows most of the protein secondary structures, but it does not resolve amino acid sidechains. To identify proteins in the stalk, neck and head regions of the RS3 cryo-EM density, we used an integrated approach, including docking of previously published and AlphaFold2^41^-predicted protein models of individual proteins or protein complexes, based on our proteomic results and COLORES-based exhaustive search of the mouse proteome^42,43^ (see method for details). This approach allowed us to completely and unambiguously localize 14 RS3 proteins (10 previously unidentified, Table 1), which are all unique to RS3 except for CFAP91, which connects RS2 and RS3. Fitting 16 RS3 subunits to all resolved RS3 cryo-EM densities resulted in an atomic model of the entire RS3 structure, plus 11 interacting proteins, such as the heavy chain of both IDA *d* (DNAH1) and IDA *g* (DNAH6), and IDA light chain DNALI1 (Fig. 2 and Supplementary Figs. 5-7; deposited as PDB: 9D2F). In addition, the highly flexible domains of two stalk-neck region proteins, CATIP and PRKAR2A, could be putatively placed (Supplementary Fig. 8), but they are not included in the deposited “high-confidence” atomic model. Most (9 to 10) or even all 14 RS3 proteins are highly conserved among ciliated eukaryotes (RS3 head, neck, and base) or animals, respectively. Our results also correct a previously published model that incorrectly fitted RS1 and RS2 proteins to a low resolution cryo-EM map of the RS3 head and neck region^18^.

**Table 1.** RS3 proteins identified by a combination of proteomic analysis of mouse WT and *AK7^-/-^* axonemes and molecular modeling into the cryo-EM structure of RS3. Protein-related diseases in humans and mice are listed.

| Protein | UniProt ID | MW (kDa) | Average ratio of abundance (AK7<sup>-/-</sup>/ WT) | Location in RS3 | Protein-related diseases |
| --- | --- | --- | --- | --- | --- |
| Newly identified RS3 proteins |  |  |  |  |  |
| AK7 | F8WIC0 | 83 | 0.04 | head-neck | <sup>a</sup>Male infertility<sup>40,93</sup>, <sup>a</sup>PCD<sup>39,94,95</sup> |
| AK9 | G3UYQ4 | 219 | 0.03 |  | <sup>a</sup>Male infertility<sup>96</sup>, <sup>c</sup>Hydrocephalus<sup>97</sup> |
| MDH1B | Q5F204 | 56 | 0.05 | neck | No disease association |
| MDH1 | P14152 | 37 | 0.63 |  | <sup>c</sup>Epilepsy<sup>98</sup> |
| <sup>d</sup>C10ORF53 | Q3KNL4 | 10 | 0.07 | stalk | No disease association |
| AKAP14 | Q3V0I7 | 58 | 0.05 |  | No disease association |
| CATIP | B9EKE5 | 59 | 0.35 |  | <sup>a</sup>Male infertility<sup>99</sup> |
| PRKAR2A | P12367 | 45 | 0.35 |  | <sup>c</sup>Fibrolamellar carcinoma-like tumors<sup>100</sup> |
| MORN5 | Q9DAI9 | 20 | 0.36 |  | <sup>c</sup>Non-syndromic cleft lip<sup>101</sup> |
| STYXL1 | Q9DAR2 | 38 | 0.50 |  | <sup>b</sup>Male infertility<sup>47</sup>, <sup>c</sup>Epilepsy<sup>48</sup> |
| Previously known RS3 proteins |  |  |  |  |  |
| LRRC23 | O35125 | 39 | 0.05 | head | <sup>a</sup>Male infertility<sup>21,22</sup> |
| CFAP91 | Q8BRC6 | 92 | 1.06 | stalk-base | <sup>a</sup>Male infertility<sup>25</sup> |
| CFAP61 | Q8CEL2 | 143 | 0.93 | base | <sup>a</sup>Male infertility<sup>24,102,103</sup> |
| CFAP251 | E9Q743 | 149 | 0.87 |  | <sup>a</sup>Male infertility<sup>23,104,105</sup> |

<sup>a</sup> Disease reported in humans and mice.
<sup>b</sup> Disease reported in mice only.
<sup>c</sup> Disease reported in humans only.
<sup>d</sup> The ratio is based on one sample repeat only, abundance is too low for quantification in another sample repeat.

**Fig. 2.**
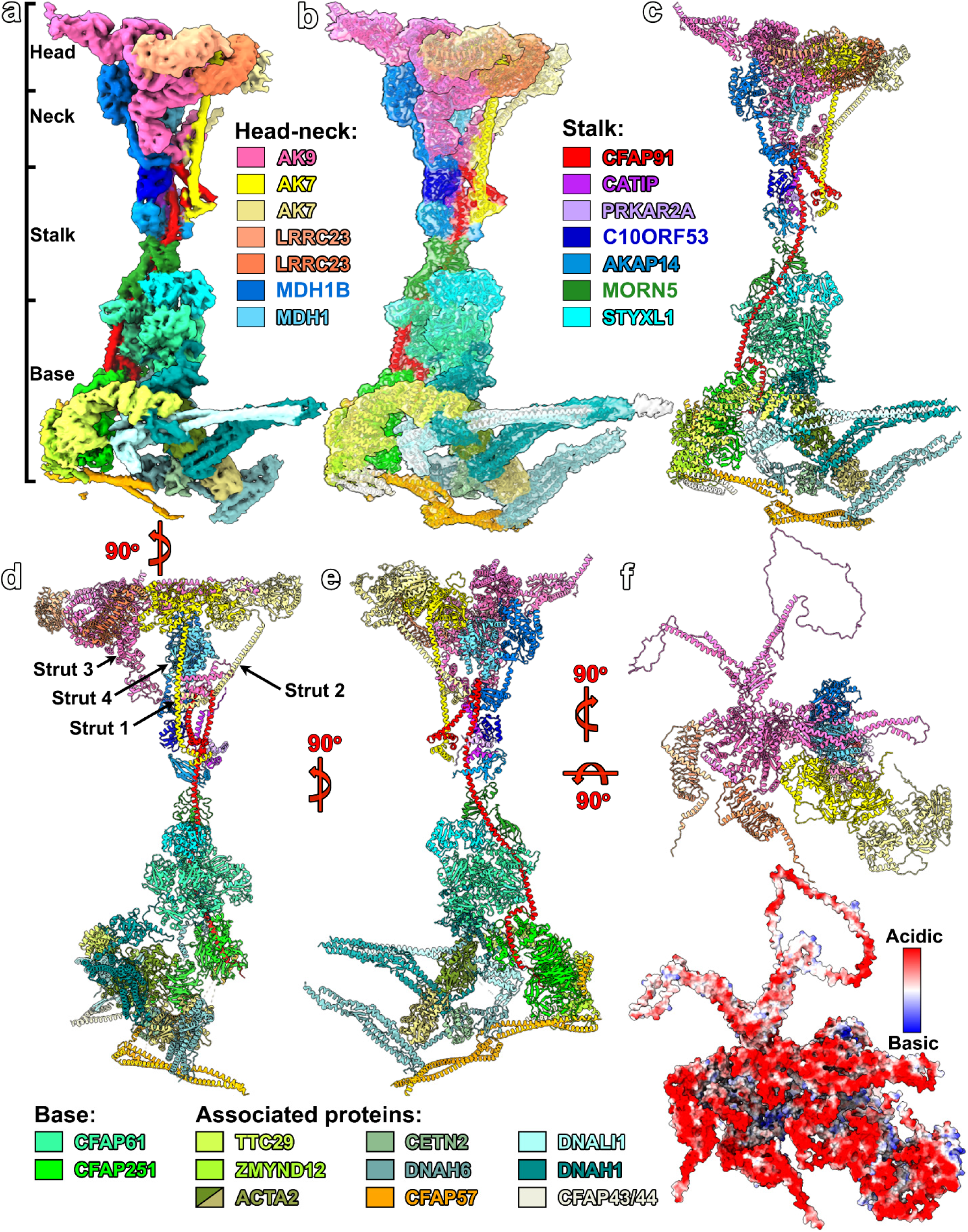
3D structure and atomic model of RS3 in mouse respiratory cilia. **a-e)** 3D reconstructed, full-length RS3 shown in cross-sectional view of the isosurface rendering colored by subunits (a), with AlphaFold2-predicted atomic models of the RS3 proteins and RS3-associated proteins (see color legends) fitted to the EM map (b), and just the atomic model seen from distal (c), front (d) and proximal (e). Note the four “struts” in the neck region (d), that are formed by long helices from the two AK7, the C-terminal domain of AK9, and the MDH1/MDH1B heterodimer to support the “flat” head region. RS3-associated proteins belong to major motor and regulatory complexes in cilia, such as the tails of IDA *g* (DNAH6) and IDA *d* (DNAH1), and dynein light intermediate chain 1 (DNALI1), IDA interacting proteins ACTA2 (2 molecules), CETN2, and ZMYND12/TTC29, the I1 dynein associated tether/tether head (T/TH) complex (CFAP43/44), and CFAP57 that links RS3 to the DMT. Additionally, CFAP91 connects the bases of RS2 and RS3. **f)** Top view of the RS3 head atomic model (top) and a charge-based surface representation showing the highly negatively charged (red) RS3 interface with the CPC projects (bottom). Note that the final refined model does not contain some of the disordered loops due to lack of cryo-EM density (a-e), but they are included in the model in panel (f) to provide a complete view of the surface potential.

### Structure of the RS3 head

The RS3 head is composed of five proteins: one AK9 (219 kDa), two AK7s (83 kDa each) and two LRRC23s (39 kDa each) (Fig. 2f and Supplementary Fig. 5). In addition to the cryo-EM map and our proteomics data supporting the presence of these proteins in the distal region of RS3 (Supplementary Fig. 2, red dots, and Table 1), LRRC23 has also been reported as RS3 component by a previous study^21^. All RS3 head proteins are rich in acidic residues, both in the ordered domains and the disordered loops. Most of the acidic residues are distributed at the largely flat surface of the RS3 head, which interfaces with the CPC projections, giving this interface a highly negative electrostatic potential (Fig. 2f, bottom), similar to the heads of RS1 and RS2^17^. This property of RS3 supports the previous hypothesis that negative electrostatic potential of RS heads helps maintain ciliary motility by preventing the radial spoke heads from colliding with or sticking to the CPC during inter-DMT sliding and axoneme bending^44^.

The RS3 head contains five catalytic AK domains (Fig. 3a-f): one in each of the two tandem-arranged AK7 molecules (Fig. 3a, e-f), and three catalytic domains in the large AK9 (Fig. 3a-d). AK enzymes catalyze the reversible transphosphorylation reaction from two molecules of ADP as substrate leading to ATP and AMP production (Fig. 3g). However, two of AK9’s three catalytic domains seem to be in an inhibited state, with the N-terminal β-hairpins of the LRRC23 molecules wedged into their substrate binding sites (Figs. 3a-c, arrowheads). Additional interactions between LRRC23 and AK9 are between the leucine-rich repeats of LRRC23 and the ATP and AMP binding loops of the catalytic domains of AK9. The AlphaFold2 complex prediction of AK9 and LRRC23 is also consistent with the observed interactions (Supplementary Fig. 6a-b). To determine biochemically if the RS3 head is important for ATP regeneration, we performed AK-activity assays on the axoneme samples from WT and FAP61-KO in *Tetrahymena,* without and with extraction (0.6 M NaCl) of RS3 from the mutant. The results show that the AK activity from FAP61-KO is significantly lower than that from WT axonemes, and especially after salt extraction the AK activity from FAP61-KO is only 50% of that from extracted WT axonemes (Fig. 3h). This is consistent with the proteomic data showing dramatic reduction of two of the four identified ciliary AKs, AK7 and AK9 from FAP61-KO in *Tetrahymena* (Supplementary Table 1 and Table 2).

**Fig. 3.**
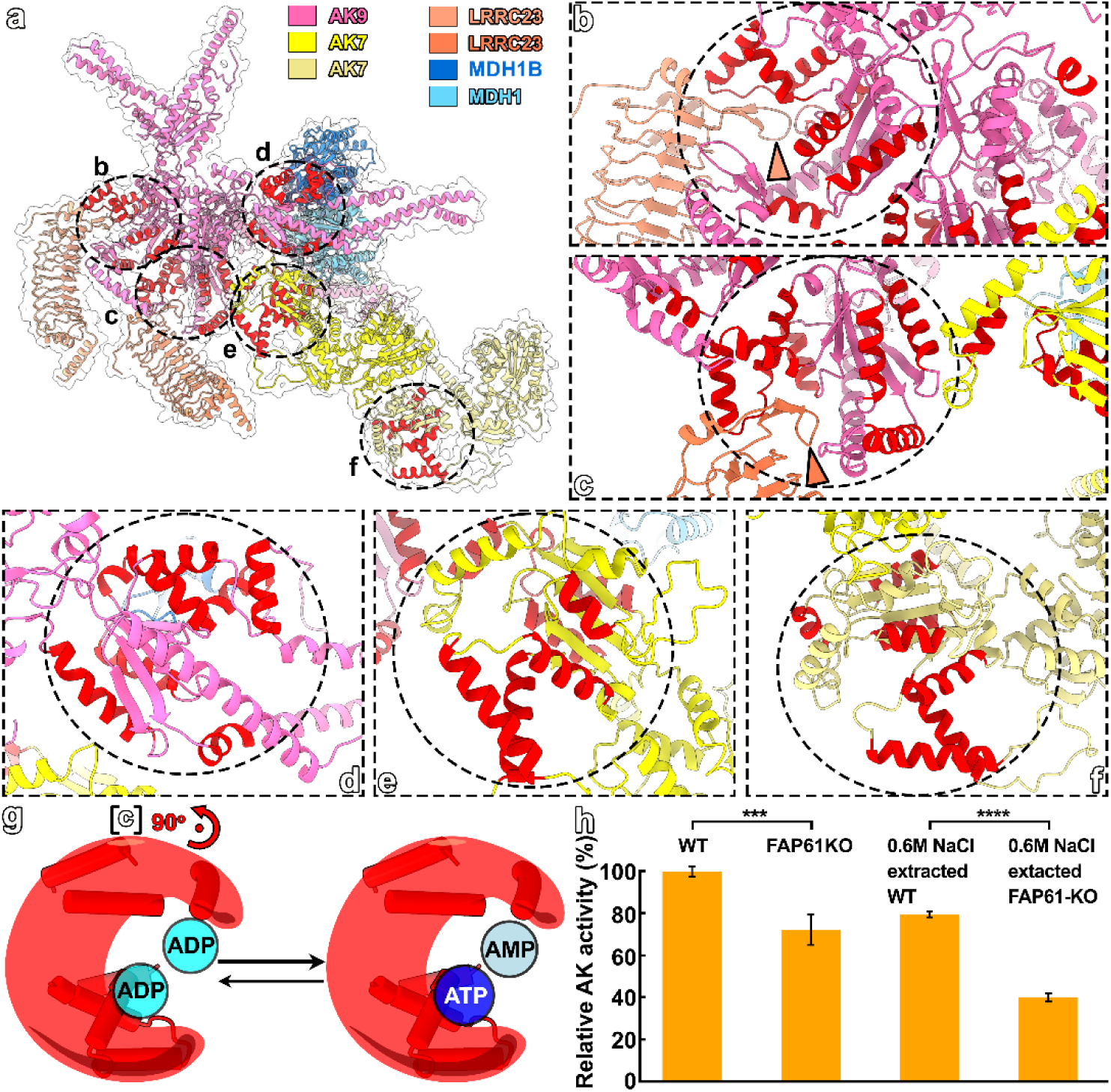
Adenylate kinase activity of RS3. **a)** Atomic model inside the transparent isosurface rendering of the RS3 head structure in top view (interface with CPC). Black circles indicate the three AK (adenylate kinase) catalytic domains of AK9 (shown in detail in b-d) and in two AK7s (shown in detail in e-f). In each domain, α-helices that are important for binding of two adenine nucleotide (2 x ADP, or ATP and AMP) are highlighted in red color. **b-c)** Zoom-ins of two AK9 catalytic domains that are blocked by the N-terminal β-hairpin (arrowheads) of two LRRC23 molecules, which interact with AK9 also through their leucin-rich repeats (a). **d-f)** Zoom-ins of another catalytic domain of AK9 (d) and of two AK7s (e-f). **g)** Graphical illustration of the catalytic domain of AK proteins (with α-helix arrangement in dark red) that catalyzes the reversible transphosphorylation reaction from two ADPs to ATP and AMP. **h)** Kinase activity assay of isolated *Tetrahymena* axonemes comparing WT and FAP61-KO with (right) and without (left) prior salt-extraction show that the AK activity of FAP61-KO is significantly lower than that from WT axonemes (*** *P* < 0.001, **** *P* < 0.0001, T test; graph shows means ± SD; n = 4 each).

### Structure of the RS3 neck

The C-terminal regions of AK7 and AK9 extend into the RS3 neck region; together with a malate dehydrogenase 1/1B (MDH1/MDH1B) heterodimer they form four angular struts that converge to a focal point at the distal end of the stalk region, where the five molecules interact with the “hook-shaped” C-terminal segment of CFAP91 (Figs. 2a-e, 4a-d. This structural framework is reminiscent of a “fixed umbrella construction” that is architecturally known for stability against external forces^45^ -- four load-bearing ribs (i.e., struts) extend from the central shaft (i.e., stalk/CFAP91) outward in a radial fashion to support a flat, canopy-like structure at the top (i.e., RS3 head). Several of the proteins involved contain extended helical segments, such as the 90 Å long C-terminal alpha-helix of both AK7 proteins that connect the head with the stalk, and the 30 nm long C-terminal segment of CFAP91, that span the RS3 base, stalk, and neck regions as a RS3 backbone (Figs. 2a-e, 4a).

Though the MDH family of enzymes is typically known to function as homodimers, MDH1/MDH1B heterodimerization in RS3 (Fig. 4a-b) is supported by two prominent features in the cryo-EM map: 1) the density for one of the MDH subunit contains a small N-terminal globular domain linked to the catalytic domain through a single helix (Fig. 4a and Supplementary Fig. 7a), which is present in MDH1B but not in other MDH family members; and 2) the density of the catalytic domain dimer shows obvious asymmetry (e.g., two corresponding helices near the pseudo-2-fold symmetry axis are straight and kinked, as predicted for MDH1B and MDH1, respectively). Our proteomic data show a significant reduction of MDH1B, but a smaller reduction of MDH1 (Supplementary Fig. 2 and Table 1), likely due to additional MDH1 molecules elsewhere in the axoneme. Supporting this, MDH1 was previously reported in proteomic analyses of WT *Chlamydomonas* axonemes^4^ that lack the RS3 stalk/neck/head regions^19^. The MDH1 isoform is known to be a key component of the Malate-Oxaloacetate shuttle that is important for maintaining the NADH/NAD^+^ ratio and cellular redox balance^46^. Though it is not known whether the MDH1/MDH1B heterodimerization affects the catalytic activity (i.e., generating NADH), it appears important for the structural integrity of RS3, because the N-terminal global domain in MDH1B serves as spacer between the neck and the stalk region by interacting with CFAP91, C10ORF53, and AK9 (Fig. 4d, Supplementary Fig. 6d and Supplementary Fig. 7a-b), yet a MDH1B homodimer would sterically clash with the RS3 head structure.

**Fig. 4.**
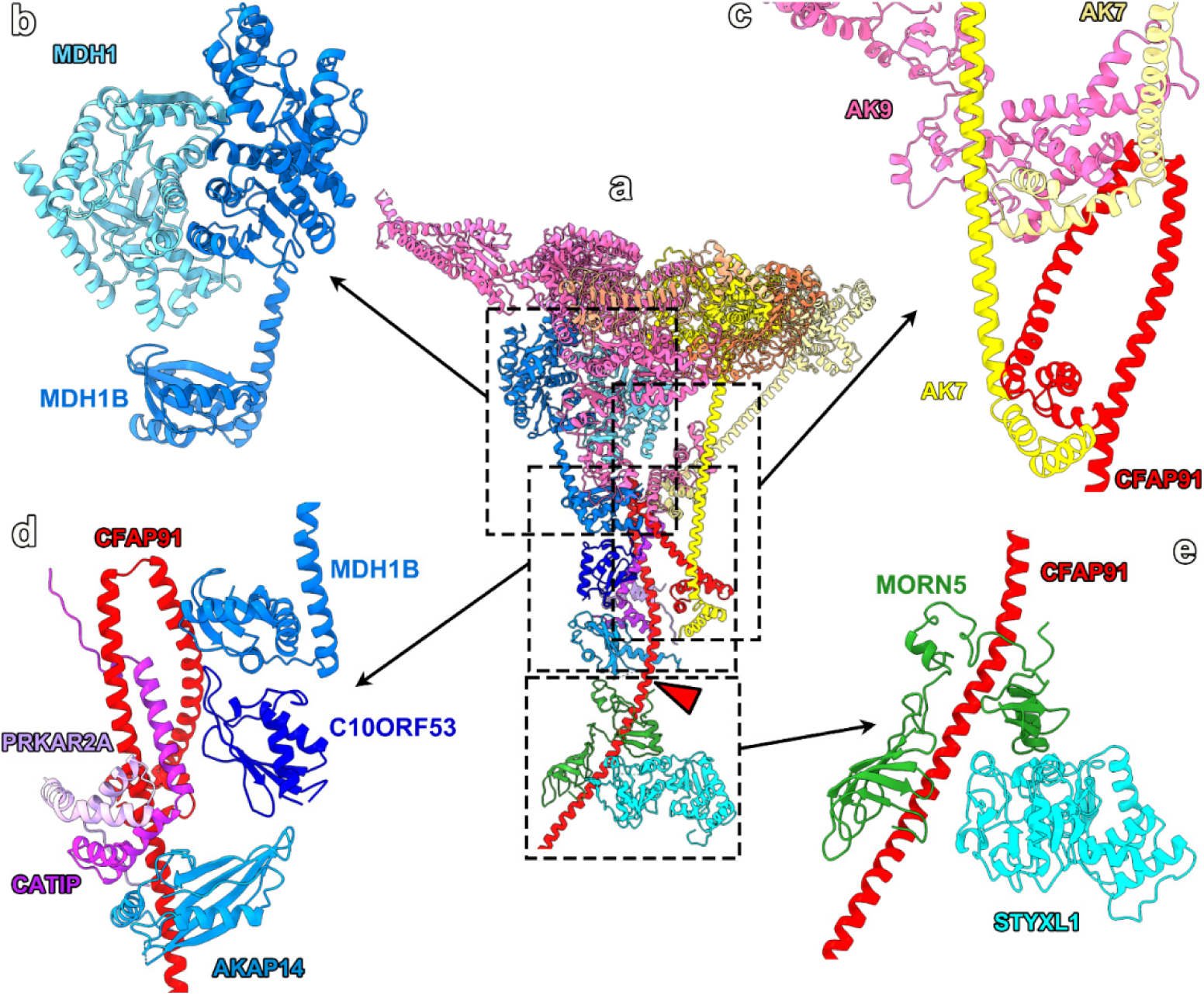
Interactions between the newly identified RS3 proteins. **a)** Atomic model of the RS3 head-neck-stalk region in cross-sectional orientation seen from distal. Boxes indicate the location of the zoom-ins shown in (b-e), which were rotated and selected subunits hidden for clarity of the region of interest. Note the “naked” region of CFAP91 (amino acids 620 to 625; arrowhead) that does not interact with other proteins. **b)** Malate dehydrogenase 1 (MDH1) and MDH1B form a heterodimer; note the extra N-terminal domain of MDH1B compared to MDH1. **c)** Tight interactions between long helical fragments of the two AK7 and the C-terminal region of CFAP91, takes nearly a 180°-turn at this region, forming a “hook-like” structure. **d)** Interaction of the scaffold protein CFAP91 with a “stack” of (neck-)stalk proteins, i.e. the N-terminal domain of MDH1B, C10ORF53 (UPF0728 protein C10ORF53 homolog), AKAP14 (A kinase (PKA) anchor protein 14), the N-terminal domain of PRKAR2A (protein kinase cAMP-dependent type II-alpha regulatory subunit) and mid-region of CATIP (ciliogenesis-associated TTC17-interacting protein). **e)** MORN repeat-containing protein 5 and STYXL1 (serine/threonine/tyrosine interacting-like 1) interact with CFAP91.

### Structure of the RS3 stalk

CFAP91 (92 kDa) has a highly extended, mostly alpha-helical structure that connects the DMT docking sites from RS2 and RS3, and then spans the entire base and stalk region of RS3 until the beginning of the neck region. Such an extended conformation of CFAP91 would be expected to be rather flexible on its own, but it is stabilized by interactions with multiple proteins in the base, stalk and neck, forming a ∼30 nm long backbone for RS3 (Figs. 2a-e, 4a,c-e and Supplementary Video 1), similar to the RSP3 (AKAP) subunit in RS1 and RS2 (Supplementary Fig. 9). The distal part of the stalk is formed by C10ORF53 (10 kDa) that interacts with the N-terminal domain of MDH1B, and the docking domains of both cAMP-dependent protein kinase type II-alpha regulatory subunit (PRKAR2A; 45 kDa) and ciliogenesis-associated TTC17-interacting protein (CATIP; 59 kDa) with a long alpha-helix that is buttressed into the hook-shaped C-terminal end of CFAP91 (Figs. 2a-e, 4d and Supplementary Fig. 7b-c and Supplementary Video 1). This is followed by the A-kinase anchoring protein 14 (AKAP14; 58 kDa) and two proteins that line the proximal part of the stalk and interact with the RS3-base protein CFAP61 (143 kDa), i.e., the MORN repeat-containing protein 5 (MORN5; 20 kDa) and Serine/threonine/tyrosine-interacting-like protein 1 (STYXL1; 38 kDa) (Figs. 2a-e, 4d,e and Supplementary Fig. 7c-d and Supplementary Video 1). Complex-formation between MORN5, STYXL1 and the helical segment formed by CFAP91 residues 580-620 is correctly predicted by AlphaFold2 (Supplementary Fig. 7d) and likely stabilizes this CFAP91 segment, providing an explanation for the previously reported importance of STYXL1 for cilia function. Mutations in STYXL1 have been associated with epilepsy disease in humans and male infertility in mice^47,48^. Between AKAP14 and MORN5 is a small “naked” CFAP91 segment (residues 620-625) that does not interact with other proteins (Figs. 2 a-e, 4a, Supplementary Fig. 7d). This slimmest part of the stalk is likely a pivot point that offers RS3 some flexibility to accommodate spatial changes between the DMTs and CPC during ciliary beating, explaining why the RS3 base/stalk and RS3 stalk/neck/head segments had to be reconstructed independently to improve the resolution (Supplementary Fig. 4a). Consistent with these protein assignments, our proteomics data show that MDH1B, C10ORF53, and AKAP14 were greatly reduced in the *AK7^-/-^* cilia (Supplementary Fig. 2, red dots, and Table 1), whereas PRKAR2A, CATIP, MORN5, STYXL1 and MDH1 were also reduced, but not as significantly (Supplementary Fig. 2, blue dots). In addition, AKAP14, PRKAR2A, CATIP, STYXL1 and MDH1B were previously reported as ciliary proteins with unknown roles^4,31,49,50^.

The secondary structure features of most RS3 proteins are well resolved in our cryo-EM map, except the functional domains of CATIP and PRKAR2A. Of these two proteins, only the small segments that attach the proteins to the RS3 stalk by interacting with CFAP91 and each other (Fig. 4d and Supplementary Fig. 7c), are clearly resolved. In contrast, the larger N-terminal and C-terminal global domains, respectively, could only be putatively placed in weak densities observed in the 3D reconstruction of the neck region of RS3 between the MDH1/MDH1B heterodimer and the C-terminal long helix of the peripheral AK7 molecule (Supplementary Fig. 8; these domains are not included in the deposited “high-confidence” atomic model of RS3). The globular domains of CATIP and PRKAR2A connect to the RS3 stalk via long, disordered linker regions that could provide both functional domains with considerable positional flexibility, with predicted gyration radii of up to 10 nm and 25 nm, respectively, around their stalk-attachment sites (Supplementary Fig. 8h-i). AKAP14 is known to recruit protein kinase A (PKA), and PRKAR2A is a regulatory subunit of PKA, suggesting that they are involved in recruiting and regulating PKA to catalyze local phosphorylation events for regulating ciliary structure and motility^49^.

## Discussion

### RS3 plays a central role in ciliary regulation

Considering the important roles of cilia for human health and fertility, and that defects in ciliary motility and assembly result in a range of “ciliopathies”^1–3^, it is important to better understanding ciliary structure, function and regulation. Cilia are highly complex and dynamic assemblies, requiring precise regulation and coordinated activity of hundreds of dynein motors to generate their characteristic, oscillatory beating. Failure of proper spatio-temporal regulation could activate the dyneins at the same or wrong time, resulting in paralyzed-stiff cilia (due to force balance) or uncoordinated (unproductive) waveforms, respectively^51^. Regulatory signals are thought to be transmitted from the CPC through the RSs to the IDAs, including the regulatory I1/f dynein, and to the CSC and N-DRC, and then further downstream to the ODAs (Fig. 5a-b)^52^. Previously described regulatory signals/mechanisms in cilia include calcium (especially CSC and CPC)^26,53^, cAMP and redox states^26^, ATP and ADP concentration^54^, phospho-regulation (e.g., I1-dynein/IC138^55^, ODA/p29^56^, N-DRC/DRC2^57^, and RSPH6A^58^), and mechanical force (e.g., between CPC and RS^27^). However, evidence for these regulatory mechanisms has so far been indirect, e.g., through biochemical and DMT-sliding assays of mutants lacking specific regulatory components; an understanding of control at the molecular level has been missing. Especially for RS3, whose molecular composition and structural organization were previously unknown, hampering our understanding of (unique) functional roles of RS3 and how the RS-CPC system transmits regulatory signals.

**Fig. 5.**
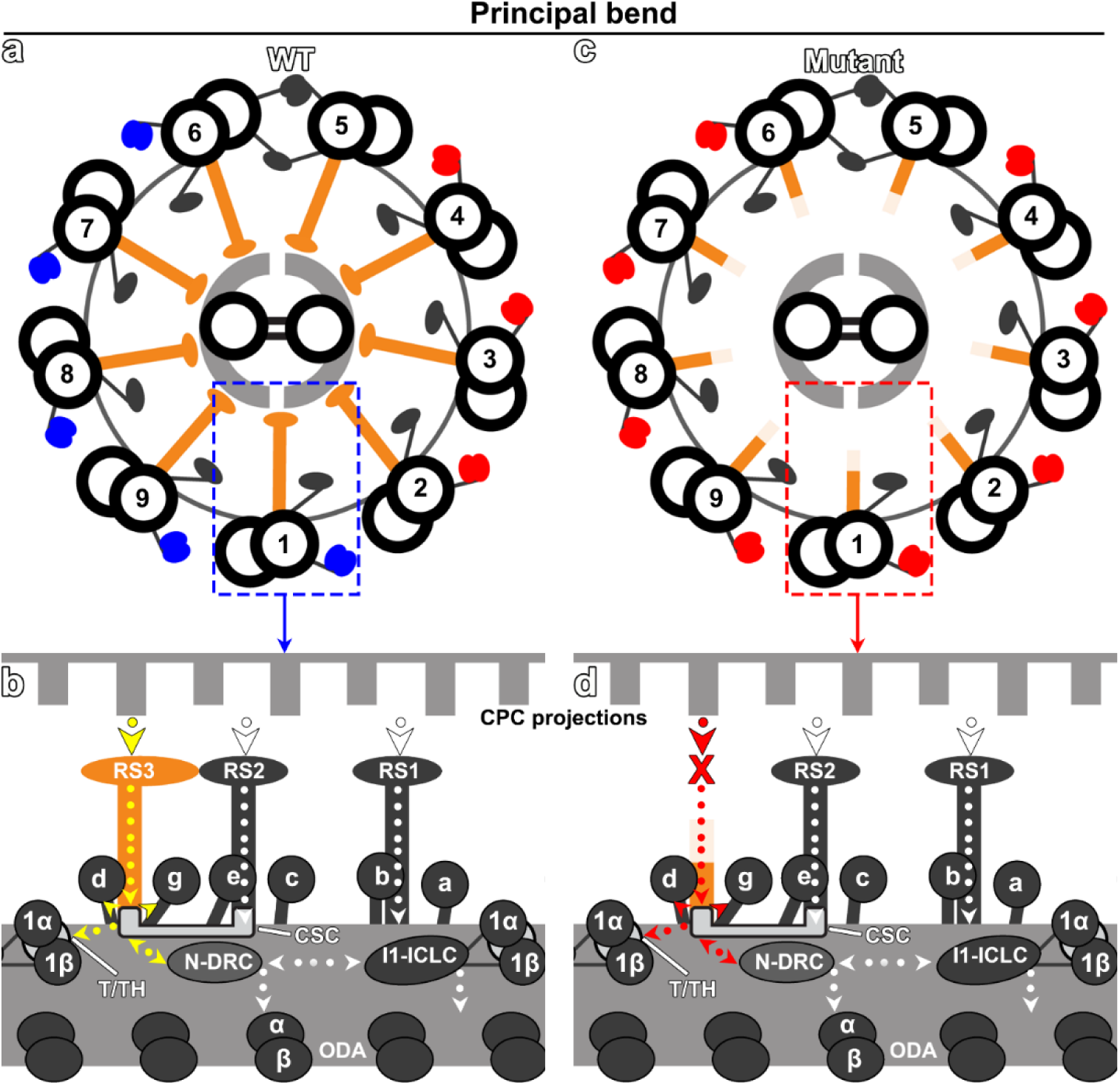
Schematic model of motility regulation in cilia. **a-d**) Cartoons show the axoneme overview in cross-section (a, c) and the 96 nm repeat in longitudinal orientation (with proximal on the right) of WT (a, b) and *AK7^-/-^* (c, d) mouse cilia. To generate ciliary bending dyneins are in their active (blue color) and inhibited states (red color), respectively, on opposing sides of the cilium (a), which switches to alter the bending direction. In WT (b), the signals are transmitted (arrows) from the CPC, through RS1 to I1 dynein (including ICLC, and associated tether/tether head, T/TH complex), through RS2 to the CSC and N-DRC, and through RS3 (yellow arrows highlight the central regulatory hub position) to the CSC, N-DRC, I1 T/TH and IDAs *d/g*. I1/f dynein and N-DRC also interact and signal further to the ODAs. In RS3 mutants (*AK7^-/-^* here), the mechanical/chemical signaling from the CPC through RS3 is disrupted (red arrows), affecting the regulation of dyneins directly and through interacting regulatory complexes.

In the *AK7^-/-^* cilia, RS3 is too short to interact with the CPC projections, thus mechano-chemical signaling through RS3 is disrupted (Fig. 5c-d). This could directly affect the regulation of RS3-associated proteins and thus major ciliary complexes, including the attached IDA motors *d* and *g*, the I1-tether/tether-head (T/TH) complex that connects to the regulatory I1/f dynein^8^, and the CSC (three of the four CSC subunits are RS3 proteins) that interacts with the N-DRC^32^ (see associated proteins in Figs. 2 and 5). Our LC-MS results from the *AK7^-/-^* axonemes also showed significant reduction of selected CPC proteins of the C1b projection (e.g., HYDIN, LRGUK, CFAP69 and SPEF2) and of the potential RS1/2 or CPC protein RSP10B^31^ (Supplementary Table 4), suggesting that the RS3 head interacts with and stabilizes axonemal assembly of these proteins. RS3 forms a central hub in an interconnected network that includes several motor and regulatory complexes in the 96 nm axonemal repeat (Fig. 5b). Therefore, it is unsurprising that mutations in 10 of the 14 RS3 proteins have previously been associated with cilia-related diseases in human or mice. As summarized in Table 1: AK7 mutations have been linked to severe PCD disease in mice (including hydrocephalus and premature death) and to human infertility (a typical symptom reported for cilia-related diseases), mutations of AK9, LRRC23, CATIP, STYXL1, all three CSC subunits (CFAP61/91/251) have been associated with male infertility in human and/or mice. In addition, MDH1 has been associated with epilepsy (a neurological disorder potentially linked to ciliary function), and MORN5 with non-syndromic cleft lip, a craniofacial abnormality related to ciliopathies.

### Phospho-regulation machinery captured

Phospho-regulation, i.e., the specific modulation of protein function and enzyme activity through their reversible phosphorylation, is a versatile and essential regulatory mechanism that coordinates a wide range of cellular processes and pathways, including metabolic and signaling pathways. Studies have revealed that phosphorylation of ODAs and IDAs is important for the regulation of beat frequency and ciliary bending^54,55^, and that the CPC-RS system regulates dynein-driven microtubule sliding by a mechanism involving protein phosphorylation^55,59,60^. For example, *in vitro* assays have shown that changes in DMT sliding velocity are mediated by phosphorylation of the IDAs^61,62^, and genetic and biochemical studies of the I1/f dynein demonstrated that the dynein intermediate chain IC138 is a key regulatory phosphorylation switch, where phosphorylated IC138 corresponds to dynein inactivation and *vice versa*^55^. Ciliary kinases and phosphatases that are thought to be involved in phospho-regulation include the serine/threonine-specific protein kinases Casein Kinase 1 (CK1) and PKA, and the serine/threonine-specific Protein Phosphatases 1 and 2A (PP1 and PP2A), which are all physically anchored to the axoneme^55,59^. CK1 and PP2A, which are required for proper regulation of microtubule sliding^63,64^, are thought to be located near the I1 intermediate-chain-light chain (ICLC) complex or the heterodimeric I1 T/TH complex^8,55^. In contrast, PKA is predicted to be anchored near the base of RS1 and RS2, as well as in the CPC based on the reported locations of two A-Kinase Anchoring Proteins (AKAPs), i.e., the RS1/2 subunit RSP3 (AKAP) and AKAP240 in the CPC, respectively^55^. AKAPs are proteins that interact with the regulatory subunits of PKA and confer sub-cellular location, and to date, AKAPs are the only known mechanism for localizing PKA in the axoneme^55^. However, previous cryo-ET or cryo-EM single particle studies have not been able to directly visualize ciliary kinase or phosphatase in the axonemal structure, except for the catalytic subunit of protein phosphatase 1 (PP1c) “buried” in two CPC projections^65^.

Here, we unambiguously located AKAP14, which is known to recruit PKA, in the RS3 stalk region where it interacts with the N-terminal domain of PRKAR2A, a regulatory subunit of PKA (Figs. 2 and 4d). The N- and C-terminal domains of PRKAR2A are connected by a long, disordered linker (72 amino acids), and we could place the C-terminal domain of PRKAR2A putatively in a blurred density in the RS3 neck region, which suggests a considerable degree of positional flexibility of this functional domain (Supplementary Fig. 8). Usually two catalytic and two regulatory PKA subunits form an inactive PKA holoenzyme. Upon cAMP-binding to the regulatory domain, the holoenzyme dissociates, releasing two free, monomeric, catalytically active subunits that then transduce the cellular signal through phosphorylation of one or more target proteins. Due to their inherently transient protein-interactions, it is notoriously difficult to visualize kinases in action. However, the presence of AKAP14 and PRKAR2A in RS3 suggests a (possibly general) mechanism, whereby an AKAP (AKAP14 here) recruits PKA regulatory subunit(s) (PRKAR2A here) to specific docking sites (RS3 stalk here), and a long, flexible linker allows the PKA regulatory subunit to sample considerable space (Supplementary Fig. 8i) to “catch” free catalytic subunit(s) to form the inactive holoenzyme. The physical anchoring of a PKA regulatory subunits to RS3, i.e., on nine DMTs and every 96 nm along each DMT, ensures that the regulatory subunits are abundant and well distributed. Together with other axonemal AKAPs, e.g., in RS1, RS2 and CPC, and recruited PKA regulatory subunits, they could cast a fine-meshed net to efficiently “catch” free, i.e., activated, PKA catalytic domains to turn off local kinase activity. Flexible attachment or release of catalytic subunits could be the reason why they are missing from our structural and proteomic data. Phospho-regulation is likely too slow to directly activate/inactivate the dyneins that drive the quick oscillatory (40 Hz) bending of cilia. Instead, phospho-regulation is likely a critical regulatory mechanism for modulating amplitude and waveform of the ciliary bend in response to cellular signals and external stimuli (e.g., capacitation of mammalian sperm flagella^58^, respiratory cilia mucus clearance^66^, backward swimming of *Paramecium*^67^, *Tetrahymena* photo-response^68^).

### RS3 is a unique cluster of metabolic proteins

A prerequisite of dynein’s motion-generating powerstroke (Fig. 6a) is the availability of ATP throughout the length of the narrow ciliary compartment, which is many micrometers long and has an only 200-300 nm wide, semipermeable opening to the cytoplasm at the ciliary base. A mouse cilium has 16 dynein heavy chains per 96 nm axonemal repeat (Fig. 1c). The ciliary bending, which switches direction every few milli-seconds, is driven by dyneins on 2-3 DMTs^51^. Thus, in a >10 µm long cilium, several thousand dyneins hydrolyze ATP and generate ADP whenever the cilium is bending. The rates of forward and reverse reactions of the dynein powerstroke cycle depend on the concentrations of substrates and products, such as ATP and ADP. Therefore, decreasing the ATP/ADP ratio along the cilia would inhibit dynein activity and impair ciliary motility. Most cilia and flagella do not contain mitochondria to support energy homeostasis, thus they likely rely on diffusion/exchange with the cytoplasm and/or local ATP generation. The latter is supported by previous experiments demonstrating that the addition of ADP (or its analogs) to isolated and washed axonemes is sufficient to generate dynein-driven motion (i.e., inter-DMT sliding)^69^. Consistent with these findings, previous proteomic studies revealed that the *Chlamydomonas* axonemal proteome (i.e., physically bound to the axoneme) contains all enzymes of the lower portion of the glycolysis pathway, including Aldolase, GAPDH, PGK, PGM, Enolase (which is also present in the cilia matrix^70^), and PK^4^. In addition, the detected MDH was thought to be involved in regenerating the NAD^+^ that is needed by the glycolytic pathway^4^. This suggests that ATP can indeed be locally produced by glycolysis to maintain a suitable ATP/ADP ratio and thus dynein motor activity along the length of the cilium^4^. However, despite of many cryo-ET and cryo-EM single particle studies of the axoneme, DMT and CPC, so far only one glycolytic protein has been localized within the axoneme, i.e., enolase in the CPC^65,71^. This has hindered a better understanding of the energy metabolism and homeostasis in cilia, which is a key prerequisite for ciliary motility.

**Fig. 6.**
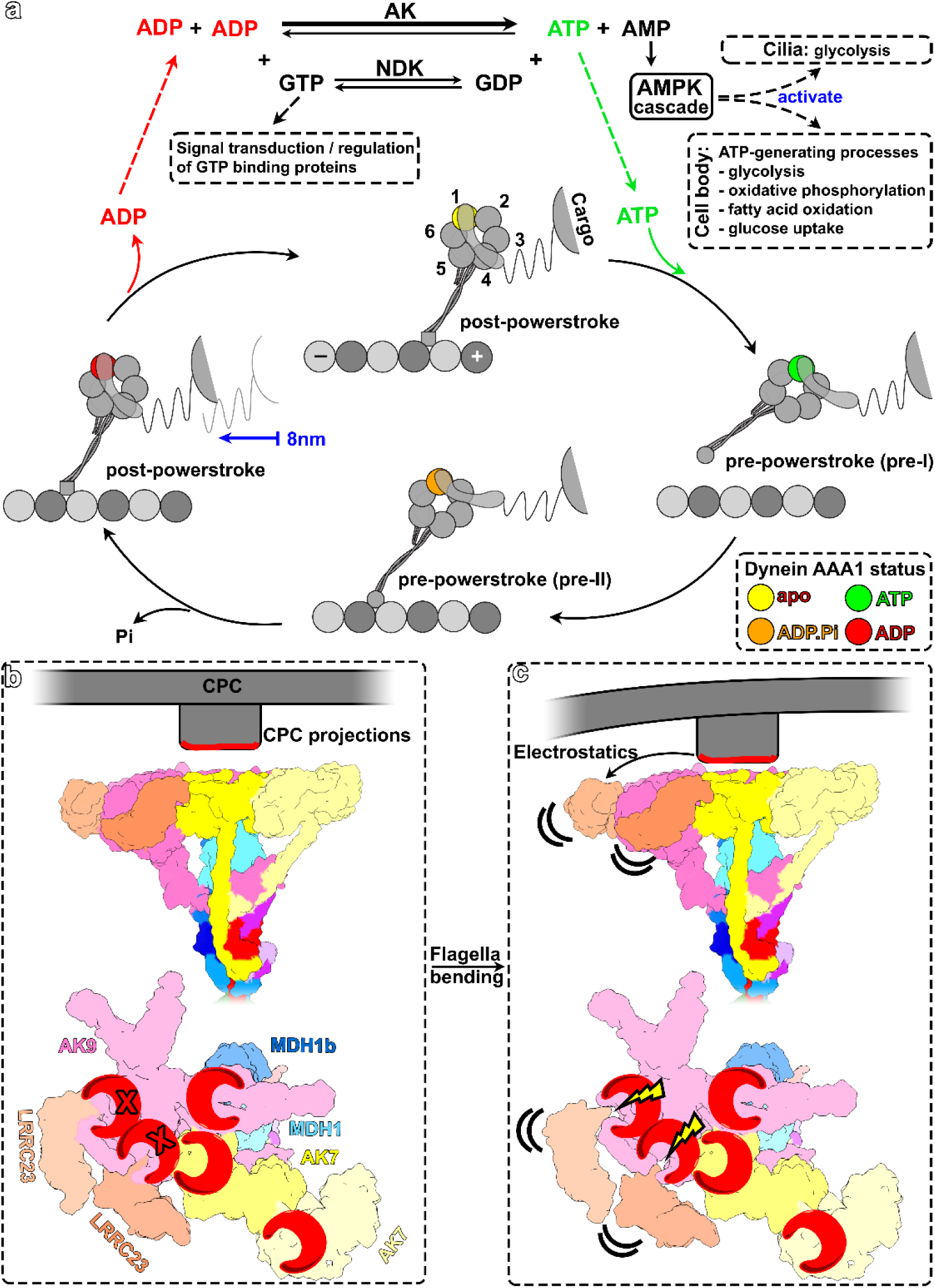
Regulatory effect of the metabolic activity of RS3 on the mechanochemical cycle of dynein. **a)** During the dynein powerstroke cycle, ATP hydrolysis drives conformational changes in the dynein motor (nucleotide state of the conformations are color coded), enabling it to transport cargo towards the microtubule minus end (-). The ATP concentration and ATP/ADP ratio affect the reaction kinetics, where high ATP and low ADP levels promote efficient motility and *vice versa*. AK enzyme activity (top), which converts 2 ADP to ATP and AMP, promotes dynein activity and thus ciliary motility by locally increasing ATP and decreasing ADP concentration. These adenosine nucleotides could affect additional pathways in cilia, e.g., through nucleoside diphosphate kinases (NDKs) or the AMPK cascade. **b, c)** Proposed molecular mechanism how RS3 could regulate ATP/ADP levels and thus dynein activity in a mechanosensitive manner: As the cilium bends, the axoneme cross-section deforms from round-to oval-shaped, bringing the CPC projections on two sides of the cilium closer to the RS heads (c). Both surfaces of the CPC/RS interface are highly negatively charged, and an increase in electrostatic force with decreasing distance could cause conformational changes, such as “pushing” the LRRC23 subunits out of two AK9 catalytic domains and increasing AK9 activity in response to mechanical force.

Here we localized two additional metabolic enzymes to RS3 (MDH1 and MDH1B) and identified three AK proteins with a total of five AK catalytic domains in the RS3 head (Figs. 2 and 3), where these metabolic enzymes also serve a structural role (e.g., as struts supporting the RS3 head), rather than being merely “associated” with the RS scaffold. The AKs likely play a key role in ciliary adenine nucleotide homeostasis by locally regenerating ATP, while reducing the ADP concentration (Fig. 6a top) and increasing AMP, which could further enhance ATP-generating processes in the cilium and cell body through the AMP-activated protein kinase (AMPK) cascade (Fig. 6a top). In addition, ciliary nucleoside diphosphate kinases (NDKs) (e.g., the highly conserved RSP23 and FAP67^28,72^) could use ATP to generation GTP and activate ciliary GTP-binding proteins that play roles in signal transduction and enzyme regulation (Fig. 6a top).

The clustering of five metabolic enzymes to a single axonemal complex, which is an efficient mechanism to ensure high abundance and regular distribution of these enzymes along the entire length of the cilium, makes RS3 unique and likely a key player in ciliary energy metabolism. The RS3 neck location might bring the MDH-heterodimer in proximity to the CPC-bound enolase, which is part of the glycolysis pathway that requires NAD^+^ regeneration to continue producing ATP. However, it is unclear where the remaining glycolytic enzymes localize in the axoneme, and whether their relative locations impact the efficiency of the glycolysis pathway.

In the isolated DMT sample used here for 3D structure determination, two of AK9’s catalytic domains appear to be inhibited by the associated LRRC23 subunits. However, in the native cilium, the highly negatively charged surface of the RS3 head would be close to the also highly negatively charged surfaces of the CPC projections (Fig. 6b). During axonemal bending, the usually round axonemal cross-section (in straight cilia) is compressed to an oval-shaped cross-section, pressing the CPC projections in parts of the axoneme closer to and even against the RS heads (Fig. 6c). This mechanical force is thought to play a role in the CPC-RS regulation of dynein activity and thus ciliary motility^27^, but the signal transduction mechanism on a molecular level is unknown. Here we propose a mechanism where the electrostatic and/or mechanical force exerted by the CPC on the RS3 head could induce conformational changes of the RS3 head proteins, such as slightly rotating LRRC23 so that the LRRC23 hairpin is pushed out of the two catalytic pockets of AK9 and the catalytic activity and ATP-regeneration by AK9 increases in a mechano-sensitive manner (Fig. 6c). This mechanism would require that RS3 can provide some resistance against the pushing force by the CPC without breaking. Consistently, a general theme of RS architecture seems to be structural stability, e.g., several struts supporting the RS-heads and a robust RS-base, combined with some pivotal flexibility at the slim stalk (Supplementary Fig. 9).

Our results suggest that RS3 not only has important regulatory hub and mechanochemical signal transduction roles, but that it also has the unique function to locally clustering proteins that mediate energy metabolism and homeostasis in cilia. Our previous cryo-ET studies of cilia from human PCD patients with mutations of the RS1 and RS2 proteins RSPH1 and RSPH4A, respectively, revealed primary RS head defects of RS1 and RS2, and severe secondary CPC abnormalities, resulting in [9+1], [9+0], [9+4], and [(8+1)+0] axonemal arrangements, whereas only 10% of the axonemes showed the typical [9+2] organization (Supplementary Fig. 10a-d). In contrast, 69% of the *AK7^-/-^* mouse cilia with PCD-causing RS3 defects showed the typical [9+2] arrangement and the remaining cilia still retained one CPC microtubule in [9+1] axonemes (31%), which is similar to isolated WT mouse cilia due to the sample preparation (compare Supplementary Fig. 10e, f). Yet phenotypically, the RS3 defects cause more severe PCD symptoms and premature death in homozygous *AK7*^-/-^ mice than PCD caused by RS1 and RS2 defects in human ciliopathy patients and mouse mutants^73–75^, underlining the distinct and critical roles of RS3 as regulatory hub and cluster of metabolic proteins in cilia. The atomic model of full-length RS3 provides unprecedented details of a unique protein network and lays the foundation for future characterization of how individual RS proteins contribute to regulating dynein activity and ciliary motility, doubtlessly impacting our understanding of ciliary diseases.

## Materials and Methods

### Animals

All animal maintenance and experimental procedures were conducted in compliance with the National Institutes of Health Guide for the Care and Use of Laboratory Animals, the institutional guidelines, and with approval from the Institutional Animal Care and Use Committee (IACUC) of the University of Texas Southwestern Medical Center, of the Children’s Hospital Boston, and of the Mayo Clinic. The *AK7^-/-^* [FVB/TH67(TRE-Hmox1-4467)] mouse strain was generated by Dr. S. Alex Mitsialis^39^ and maintained on a FVB background. Heterozygous *AK7^+/-^* mice were provided by Dr. S. Alex Mitsialis for breeding in the animal resource center (ARC) UT Southwestern Medical Center.

### WT and *AK7^-/-^* mice breeding, maintenance, and genotyping

Mice used for this study were about 4 weeks old for proteomics (comparing WT and mutant), and up to 2 months old for cryo-EM experiments (WT only). The genotype of mice was confirmed at day 12-17 by PCR using three primers (comF: GACATCCTGCAGCAAATAGATAG, wtR: TTCCCCATGGCTTGAGATC, mutR: GACCGTTCAGCTGCAGATTA). Homozygote *AK7^-/-^* mice have primary ciliary dyskinesia (PCD) disease with reduced viability, growth retardation and hydrocephalus. Therefore, animals were closely monitored. Analgesia treatment and other procedures were performed in accordance with our IACUC-approved protocols, including treating animals with 0.5 mg/kg sustained release Buprenorphine (Bup-SR) every 48-72 hours to affect pups older than 14 days for analgesia; pups with severe phenotype less than 14 days of age were either euthanized or treated with analgesia based on ARC veterinary consulting.

### Axoneme preparation from *Tetrahymena*

*Tetrahymena thermophila* WT (CU428) and FAP61-KO^7^ cells were grown to a density of approximately 3×10^5^/ml in 1 L of super proteose peptone (SPP) medium (1% proteose peptone, 0.1% yeast extract, and 0.2% glucose) at room temperature. The axonemes were isolated as previously described^76^ with minor modifications. Briefly, cells were collected by centrifugation, washed with 10 mM Tris pH 7.5, and resuspended in 20 ml of 10 mM Tris, 50 mM sucrose, 10 mM CaCl_2_, pH 7.5. Cilia were detached from the cells by adding 350 µl of 0.5 M acetic acid. After 1 min, 180 µl of 1 M KOH were added, followed by adding protein inhibitors, such as aprotinin and PMSF. The cilia (supernatant) were separated from the cell bodies and debris (pellet) by centrifugation at 1,860 g for 5 min twice. Then the cilia were collected by centrifugation at 10,000 g for 15 min, and demembranated in HMEEK buffer (30 mM HEPES pH 7.4, 1 mM EGTA, 5 mM MgSO4, 0.1 mM EDTA, 25 mM KCl) with 1% IGLPAL 630 for 30 min. The axonemes were collected by centrifugation at 10,000 g for 10 min. The pelleted axonemes were then gently resuspended in either HMEEK buffer (un-extracted sample) or a HMEEK buffer containing 0.6 M NaCl (extracted sample). Both samples were incubated on ice for 30 min, re-pelleted by centrifugation at 10,000 g for 10 min, resuspended in HMEEK buffer, and kept on ice until use.

### Axoneme and DMT preparation from mice

WT, *AK7^-/-^* and *AK1^-/-^* mice 2.5-4 weeks old were anesthetized using ketamine/xylazine IP to immobilize them. A ventral midline incision (2-3 cm long) was made from chin to the front of the sternum, the trachea removed and placed in a 15-ml tube with sterile saline solution (0.9% NaCl) buffer at 4°C. For cryo-ET samples, this step was performed at the Children’s Hospital in Boston (*AK7^-/-^* and WT) and at the Mayo Clinic (*AK1^-/-^*), respectively, before shipping the tracheas overnight to the University of Texas Southwestern Medical Center. The tracheal tubes were carefully sliced open, twice washed, and centrifuged at 300*g* for 2 min at 4°C using saline solution. The axoneme isolation was performed as previously described^19^ with some optimizations. Briefly, after removing the saline solution, 0.5 ml of ice cold deciliation buffer (20 mM Tris pH 7.5, 50 mM NaCl, 10 mM CaCl_2_, 1 mM EDTA, 0.1% Triton X-100, 7 mM β-mercaptoethanol, 1% protease inhibitor cocktail (Sigma P8340)) was gently added to the ciliated inner surface of the trachea. The tubes with the samples were gently rotated for 2 min, then the supernatant containing the cilia was transferred to a 1.5 ml microcentrifuge tube. This process was repeated two times with the tracheal tissue to ensure that the majority of cilia were dissociated from the respiratory epithelium. Mucus and cellular debris were gently pelleted by centrifugation at 500*g* for 1 min. The supernatant was further centrifuged at 10,000*g* for 5 min to collect the demembranated axonemes.

For mass spectrometry (WT and *AK7^-/-^*), the axoneme pellet was gently resuspended in HMEEK buffer (30 mM HEPES pH 7.4, 25 mM KCl, 5 mM MgSO_4_, 1 mM EGTA, 0.1 mM EDTA) and kept at -80°C until further analysis. For cryo-ET (WT, *AK7^-/-^* and *AK1^-/-^*), the axoneme pellet was also gently resuspended in HMEEK buffer, but then immediately used for cryo-sample preparation. For the preparation of DMT for cryo-EM single particle analysis from 2 months old WT mice, the axoneme pellet was gently resuspended in HMEEK buffer containing 1% protease inhibitor cocktail (Sigma P8340). Axonemes were splayed into DMTs by treating them with 1 mM ATP and 0.5 mM NaCl in HMEEK at room temperature for 15 min while gently rotating the tubes every 5 min. Splayed axonemes were centrifuged at 15,000*g* for 30 min at 4°C. The pellet of DMTs was gently resuspended in HMEEK and immediately used cryo-sample preparation.

### Liquid chromatography–mass spectrometry and quantitative analysis

For *Tetrahymena*, pellets of WT and FAP61-KO axonemes - with or without high-salt extraction - were dissolved in a lysis buffer (7 M urea, 2 M thiourea, 4% 3-[(3-cholamidopropyl) dimethylammonio]-1-propanesulfonate, 65 mM dithiothreitol, and 2% IPG buffer, pH 3-10 nonlinear; GE Healthcare) with vigorous stirring for 30 min. Then the sample was centrifuged at 45,000 × g for 1 h to remove insoluble components. The supernatant was stored at -80 °C. Protein concentrations were determined using a 2-D Quant Kit (GE Healthcare). A total of 35 µg of proteins for each sample was separated on NuPAGE 4-12% Bis-Tris Mini Gels (Novex, Life Technologies, Grand Island, NY). The gels were fixed and stained with Coomassie blue G-250. Each sample lane was cut into four pieces that where then separately washed in 50% acetonitrile and analyzed by micro capillary reverse-phase HPLC nano-electrospray tandem mass spectrometry on a Thermo LTQ-Orbitrap mass spectrometer at the Harvard Microchemistry and Proteomics Analysis Facility, Harvard University.

For mice, pellets of WT and *AK7^-/-^* mouse axonemes were stored at -80°C until sufficient material is collected for mass spectrometry (i.e., from 20-30 WT or 60-70 AK7^-/-^ mice). Then the samples were dissolved in Laemmli buffer (Bio-Rad) with 357.5 mM β-mercaptoethanol and heated in 98°C for 5 min. A total of 30 µg of proteins for each sample was separated on a 4-12% gradient SDS-polyacrylamide gel (Genscript Biotech). After the dye-front migrated on the gel for 2.5-3.0 cm, the gel was fixed and stained with Coomassie Brilliant Blue for 30 min and destained until the background was clear. Each sample lane was cut into four pieces that were then separately processed. After dicing the sample into about 1 mm^3^ cubes, in-gel trypsin digestion, liquid chromatography-mass spectrometry (LC-MS) and peptide identification were conducted by the proteomics core facility at the University of Texas Southwestern Medical Center. Briefly, samples were digested overnight with trypsin (Pierce) following reduction and alkylation with DTT and iodoacetamide (Sigma-Aldrich). The samples then underwent solid-phase extraction cleanup with an Oasis HLB plate (Waters), and the resulting samples were injected onto an Orbitrap Fusion Lumos mass spectrometer coupled to an Ultimate 3000 RSLC-Nano liquid chromatography system. Samples were injected onto a 75 μm i.d., 75-cm long EasySpray column (Thermo) and eluted with a gradient from 0–28% buffer B over 90 min. Buffer A contained 2% (v/v) acetonitrile and 0.1% formic acid in water, and buffer B contained 80% (v/v) acetonitrile, 10% (v/v) trifluoroethanol, and 0.1% formic acid in water. The mass spectrometer operated in positive ion mode with a source voltage of 1.5-2.5 kV and an ion transfer tube temperature of 275 °C. MS scans were acquired at 120,000 resolution in the Orbitrap, and up to 10 MS/MS spectra were obtained in the ion trap for each full spectrum acquired using higher-energy collisional dissociation (HCD) for ions with charges 2–7. Dynamic exclusion was set for 25 s after an ion was selected for fragmentation. Raw MS data files were analyzed using Proteome Discoverer v3.0 (Thermo), with peptide identification performed using Sequest HT searching against the mouse protein database from UniProt. Fragment and precursor tolerances of 10 ppm and 0.6 Da were specified, and three missed cleavages were allowed. Carbamidomethylation of Cys was set as a fixed modification, with oxidation of Met as a variable modification. The false-discovery rate (FDR) cutoff was 1% for all peptides. Peptide abundances are defined as the peak intensity of the most abundant charge state for the peptide ion. Three independent datasets (each including WT and *AK7^-/-^*) were combined and sorted based on normalized abundance using the *AK7^-/-^*comparison to WT. To ensure consistency across datasets, internal normalization was performed using the average of N-DRC and axonemal ruler proteins for each dataset. Sorting criteria were based on the average *AK7^-/-^* comparison to WT ratio. A volcano plot was generated by plotting -log10 p-values against log2Fold change using GraphPad Prism (version 10; GraphPad Software, Inc, La Jolla, CA), which served as a guideline for selecting RS3 protein candidates.

### Adenylate Kinase Assay

Adenylate kinase activity was measured as previously reported^77^. Briefly, 20 µg of isolated *Tetrahymena* axonemes were added to a buffer containing 70 mM glycylglycine, pH 8.0, 10 mM glucose, 5 mM MgSO_4_, 1 mM NADP, 5 units/ml hexokinase, and 1 unit/ml glucose-6-phosphate dehydrogenase in a cuvette to a volume of 0.5 ml with 10 µM dynein inhibitor sodium orthovanadate^78^. After the cuvette was preincubated for 5 minutes at room temperature, the reaction was started by adding 10 µl of 0.1 M ADP to a final concentration of 2 mM. Adenylate kinase activity was calculated by measuring the increase in absorbance at 340 nm induced by the production of NADPH.

### Negative staining EM

400-mesh copper grids were carbon coated in house and glow discharged (30 s at 15 mA). 3.5 µl of salt-extracted (0.6 M NaCl) axonemes from *Tetrahymena* WT and FAP61-KO were applied to the carbon side of the EM grid. Quickly, the liquid was blotted from the grid using filter paper and the grid was then floated on a drop of 2% aqueous uranyl acetate, blotted and washed using filter paper for three times, and finally air dried. Then the grid was transferred into a Morgagni electron microscope (Thermo Fisher Scientific). Images were acquired at 80 kV using a 1k x 1k CCD camera (AMT).

### Cryo-sample preparation

5-fold concentrated 10 nm colloidal gold (Sigma-Aldrich, St. Louis, MO) was coated with BSA as previously described for use as fiducial marker in cryo-ET reconstructions^79^. Holey carbon EM grids (Cu, mesh 200, R2/1 or R2/2, Quantifoil Micro Tools GmbH, Jena, Germany) were glow discharged (30 s at 30 mA) and loaded in a home-made plunge freezer. For cryo-ET imaging, 3 µl of isolated axonemes from mouse respiratory cilia and 1 µl of the BSA-coated colloidal gold solution were added to the EM grid, whereas for cryo-EM single particle imaging, 4 µl of the DMT sample were added to the EM grid. After removing excess liquid by blotting the grid from the backside with Whatman filter paper (grade 1) for 2-3 s, the grid was immediately plunge-frozen in liquid ethane. The vitrified samples were then stored in liquid nitrogen until data collection.

### Cryo-ET and Cry-EM data acquisition

Cryo samples for cryo-ET were transferred into a Tecnai F30 transmission electron microscope (Thermo Fisher Scientific, Waltham) with a side-entry cryo-holder (Gatan, Pleasanton, CA) and always kept below the devitrification temperature at the Brandeis University Cryo-Electron Microscopy Facility. The microscope was equipped with a postcolumn energy filter (Gatan) and was operated at 300 keV. Samples were imaged at -8 µm defocus, under low-dose conditions and in zero-loss mode of the energy filter (20 eV slit width). All images were recorded on a 2k ×2k charge-coupled device camera (Megascan 795; Gatan) at a magnification of 13,500×, resulting in a pixel size of 10.77 Å. Tilt-series images were recorded automatically from -65° to +65° with 2 angular increments, using the SerialEM image acquisition software^80^. The accumulative electron dose of each tilt series was limited to ∼100 e/Å^2^.

For the cryo-EM single particle data acquisition, grids were loaded into a Titan Krios transmission electron microscope (Thermo Fisher Scientific) operated at 300 kV at the UT Southwestern Medical Center Cryo-Electron Microscopy Facility. The microscope was equipped with a K3 Summit direct electron detector (Gatan) and a Bioquantum post-column energy filter (Gatan) that was operated in zero-loss mode with 20 eV slit width. A total of 50,000 movies were recorded at a magnification of 81,000×, corresponding to a calibrated pixel size of 1.079 Å and with a defocus range of -1 to -3 μm. Images were dose-fractionated into 70 movie frames with a total exposure time of 4.2 s, a dose rate of 14 electrons/pixel/second, and up to a total dose of 60 e/Å^2^. Data collection was semi-automatic utilized the SerialEM software^80^.

### Image Processing

3D tomograms were reconstructed from the tilt series using the IMOD software package^81^ with fiducial marker alignment and weighted back projection. In improve the signal-to-noise ration and thus resolution, 3D subvolumes including the 96-nm axonemal repeats were extracted from the tomograms, aligned, and averaged in 3D using the PEET software^5^ with missing wedge compensation. To further analyze structural defects of RS3 in the FAP61-KO, classification analyses and class averaging were performed on the aligned subtomograms using the PEET program^82^ with a mask to focus the analyses on RS3. The numbers of tomograms, subtomograms analyzed and the resolutions of the resulting averages are summarized in Supplementary Table 3. The resolution was estimated at the center of the DMT of the axonemal repeat using the Fourier shell correlation method with a criterion of 0.5 (Supplementary Table 3). 2D tomographic slices and 3D isosurface renderings were obtained to show the structures using IMOD^81^ and UCSF Chimera^83^, respectively.

For the single particle analysis on RS3, a flowchart of the data processing workflow can be seen in Supplementary Figures 3 and 4. Briefly, the dose-fractionated image stacks were aligned and dose-weighted using cryoSPARC^84^, and the parameters of the contrast transfer function (CTF) were estimated. A total of 280,000 particles containing RS3 were manually picked (using the conserved spacing of RSs as guide) centering on the A-tubule of the DMT. Following extraction with a box size of 650 pixels to include the DMT cross-section and the RS3 base, we performed 2D classification and excluded bad classes, resulting in a total of 273,141 particles. 3D reconstruction of the DMT was performed using the 3D atomic model of the DMT from the bovine trachea (7RRO) that was transferred into a density map using ChimeraX^85^ with a box size 650 voxels. Heterogeneous refinement was used to reconstruct the *ab initio* model. Homogenous refinement and non-uniform refinement^86^ yielded a 3D reconstruction of the mouse respiratory DMT with an overall resolution of 4-6 Å (Supplementary Fig. 3f), but the density of the RS3 was blurred.

To improve the alignment and resolution of RS3 we performed signal subtraction of the DMT from the micrographs using cryoSPARC, because the DMT dominates the alignment, and exported the particles to Relion that includes the Blush refinement algorithm^87^. A total of 191,448 particles were binned by a factor of 2 (increasing the pixel size from 1.079 Å to 2.158 Å) and reduced to a box size of 360 to lower the memory usage (Supplementary Fig. 4a). One round of 3D refinement using the Blush algorithm led to an initial 3D reconstruction of the RS3 base with a resolution of 6.5 Å, resolved most of the protein features in the base region of RS3, whereas the density for the stalk, neck, and head regions of RS3 remained poorly resolved. Therefore, two overlapping masks that covered the base-stalk and neck-head regions, respectively, were generated for additional signal subtraction and focused alignment refinement. 3D reconstruction of the neck-head region after the base-stalk signal subtraction led to a map with 8.4 Å resolution, in which secondary structural features were clearly resolved. After 3D classification of the particles one class with 37,453 particles showing strong density and the 3D reconstruction of this class with Blush refinement generated a map at 7.1 Å resolution. The refined map was B-factor sharpened using the postprocess module in Relion. A similar procedure was used to 3D reconstruct the base-stalk region, except here an additional CTF refinement improved the resolution of the final map to 4.7 Å where many amino acid sidechains were visible. The two separate 3D reconstructions were aligned based on their overlapping region and combined into one composite 3D map by taking the maximal value of each voxel from the two maps using ChimeraX^85^. Focusing on the importance of protein identification in the neck-head region, we resampled the base-stalk map (slightly reducing the quality of the map due to interpolation) to fit the neck-head map (resolution preserved) during the composite map generation, followed by protein identification and atomic model building. Resolutions of the maps were calculated according to the gold-stand Fourier Shell Correlation (FSC) criteria with the 0.143 threshold (Supplementary Fig. 4b).

### Identification of proteins in RS3 and atomic model building

Based on our cryo-ET and classification results of *AK7^-/-^* axonemes, we expected that the RS3 head and neck proteins would be significantly reduced in our mass-spectrometry data of the mutant, whereas RS3 stalk proteins would likely be less severely reduced and RS3 base proteins unaffected. Accordingly, docking of the AlphaFold2 models of the top 10 proteomic hits (i.e., proteins reduced at least 90 %) to the cryo-EM map using ChimeraX^85^ led to the assignments for all five RS3 head proteins, i.e., two copies of adenylate kinase 7 (AK7, Uniprot ID: F8WIC0), adenylate kinase 9 (AK9, Uniprot ID: G3UYQ4), and two copies of leucine-rich repeat-containing protein 23 (LRRC23, Uniprot ID: O35125). For AK9, docking of the AlphaFold2 model into the map using ChimeraX showed an excellent matching score except for the C-terminal domain. However, after repositioning the domain orientation rigidly into our cryo-EM map, the AlphaFold2 predicted secondary structure elements and overall fold of the AK9’s C-terminal domain matched well. This assignment of the C-terminal domain was further supported by AlphaFold2 complex prediction with the N-terminal domain of MDH1B (Supplementary Fig. 6d). The domains were fitted into the cryo-EM density in ChimeraX and refined with ISOLDE^88^ and Phenix^89^.

To identify the remaining RS3 neck and stalk proteins, the map was segmented into smaller parts that likely contain one or a few proteins for unbiased matching with 21,615 AlphaFold2-predicted protein models of the mouse proteome^41^, using the program COLORES as described previously^42,43^. As pointed out by Chen *et al*.^43^, the correlation coefficient scores calculated by COLORES are often unrealistically high and not reliable for ranking the candidates, especially for large proteins that occupy most of the volume of the map. Instead, we calculated the FSC between the map and the model docked into the map by COLORES using the EMAN2 package^90^. Two thresholds, 0.5 and 0.2, were chosen to report the resolutions according to the FSC curves between the map and model. Highly ranked solutions based on this metric were inspected in ChimeraX, and only those showing good fit between the maps and models were chosen for further analyses. This optimized workflow led to the identification of the following four RS3 neck-stalk proteins: malate dehydrogenase 1 (MDH1, Uniprot ID: P14152), UPF0728 protein C10ORF53 homolog (C10ORF53, Uniprot ID: Q3KNL4), A kinase (PRKA) anchor protein 14 (AKAP14, Uniprot ID: Q3V0I7), and serine/threonine/tyrosine interacting-like 1 (STYXL1, Uniprot ID: Q9DAR2) protein. The assignment of these proteins was further corroborated by AlphaFold2 multimer predictions of complexes of proteins that are placed next to one another in the map^91^. As shown in Supplementary Figures 6 and 7, the AF-predicted models of these complexes are highly similar to the corresponding maps and final refined models, both in terms of the structure of the individual proteins and of the binding mode between the proteins.

After the previous nine proteins were fit into the distal part of the RS3 cryo-EM map and the model fitting of the previously published RS3 base model with 3 RS3 proteins, the only unidentified region of the cryo-EM map was the middle of the RS3 stalk, where the lower resolution of the map did not allow COLORES to identify the proteins. However, the density clearly shows that the CFAP91 forms a long helix that spans this region, as speculated previously^18^. Therefore, we used AlphaFold2 multimer prediction to predict complexes between this segment of CFAP91 and fragments of proteins from our mass spectrometry list, whereby we incrementally decreased the threshold stringency (meaning we started with proteins up to ratio<0.2, then <0.3 and so on), and tested for predicted complex formation with a high confidence score^91^. The predicted complexes were filtered by the it scores calculated by ColabFold^91^, and then chosen if the complexes showed good fit with the density map. This strategy led to the identification of three additional RS3 stalk proteins at the reduction ratio of <0.4, i.e., MORN repeat-containing protein 5 (MORN5, Uniprot ID: Q9DAI9), cAMP-dependent protein kinase type II-alpha regulatory subunit (PRKAR2A, Uniprot ID: P12367) and ciliogenesis-associated TTC17-interacting protein (CATIP, Uniprot ID: B9EKE5).

Model building was started by fitting the previously published human RS3 base atomic model (PDB: 8J07) and the AlphaFold2 models of the newly identified proteins to the composite map of the full-length RS3. The human proteins were mutated into mouse protein sequences in ChimeraX^85^. Flexible loops without density were removed. Detailed fitting and refinement of the individual residues into the density was carried out using the ISOLDE tool as a plug-in in ChimeraX^88^. Considering the map resolution, all refinement simulations were run with secondary structure restraints. The model output by ISOLDE was further refined for coordinates and b-factors with the real-space refinement module in the Phenix package, with secondary structure and Ramachandran restraints^89^. Model validation was run with the Molprobity program as a part of the Phenix package^92^. The resolution of the model was assessed by the FSC between the model and the map (Supplementary Fig. 4b, threshold 0.5). The model statistics are summarized in Supplementary Table 3.

## Acknowledgements

We thank Dr. Chen Xu for providing EM training and management of the electron microscopes in the Louise Mashal Gabbay Cellular Visualization Facility at Brandeis University. We thank Dr. Daniel Stoddard and Jose Martinez Diaz for providing EM training and management of the UT Southwestern Medical Center cryo-electron microscope facility (funded in part by Cancer Prevention and Research Institute of Texas (CPRIT) Core Facility Awards RP170644 and RP220582). We thank Drs. David Mastronarde and John Heumann (University of Colorado at Boulder) for continued development of image processing tools, including PEET classification. This research benefitted from the computational resources provided by the BioHPC (high-performance computing) facility located in the Lyda Hill Department of Bioinformatics at UT Southwestern Medical Center. Mass spectroscopy was conducted in the UT Southwestern Proteomics Core Facility. We thank Drs. Mary Porter (University of Minnesota), Stephen King (University of Connecticut), J. Richard McIntosh (University of Colorado), Benjamin Tu (UT Southwestern Medical Center) and Justine Pinskey (University of Massachusetts) for their critical review and helpful feedback on the manuscript. This work was supported by funding by the following National Institutes of Health (NIH) and the CPRIT grants: R01GM083122 and RR140082 to D.N., and R35GM130289 (to X.Z.).

## Author Contributions

Y.Z. performed sample preparation, data collection, data processing for the cryo-EM study of mouse, sample preparation and analysis for the LC-MS of mouse, figure, and movie preparation. K.S. performed sample preparation, data collection, data processing for the cryo-ET study of mouse, sample preparation and analysis for the LC-MS and enzyme activity assay of *Tetrahymena*. L.G. performed data collection for the cryo-EM study of mouse. A.T.T. performed data analysis for the LC-MS of mouse. A.F.G., S.Z., P.P.D. and S.A.M. generated and provided the *AK7^-/-^* and *AK1^-/-^*mouse strains. X.Z. performed data processing and modeling for the cryo-EM study of mouse. Y.Z., X.Z. and D.N. wrote the manuscript. D.N. conceived and directed the study.

## Data availability

The 3D structures of the subtomogram averaged 96 nm axonemal repeats of different mouse strains have been deposited in the Electron Microscopy Data Bank (EMDB; https://www.ebi.ac.uk/pdbe/emdb/) under accession codes EMD-46484 (WT), EMD-46486 (*AK7^-/-^*), and EMD-46485 (*AK1^-/-^*). The composite cryo-EM density map and atomic coordinate of RS3 from mouse respiratory cilia have been deposited in the Electron Microscopy Data Bank and Worldwide Protein Data Bank (PDB; https://www.rcsb.org/), respectively, under accession number EMD-46494 and PDB: 9D2F.

## Code availability

The Python scripts for running the COLORES-based search, EMAN2-based FSC resolution calculation and ranking of the models are deposited into the following GitHub site: https://github.com/xuewuzhang-UTSW/Exhaustive-search-for-proteins-in-cryo-EM-map.

## Materials & Correspondence

Correspondence to Daniela Nicastro and Xuewu Zhang

